# easyClock: A User-Friendly Desktop Application for Circadian Rhythm Analysis and Visualization

**DOI:** 10.1101/2025.07.08.663749

**Authors:** Binbin Wu, William W. Ja

## Abstract

Circadian rhythms regulate a wide range of biological processes, and their precise characterization is essential for understanding behavioral and physiological fluctuations. However, existing tools to analyze circadian data often require coding expertise or rely on specific data acquisition software, limiting their general applicability. Here, we present easyClock, an intuitive and interactive application designed to streamline circadian rhythm analysis and visualization. The easyClock application enables simultaneous processing of multiple files, allowing users to preview, batch-analyze, and plot diverse sets of time series generated from high-throughput experiments. To enhance data analysis efficiency and provide comparable results, this application integrates four different analysis methods for handling data with various waveforms. Results are easily viewed and exported for any selected range of data. This free-to-use application is open-source and can be deployed on macOS computers by users without coding experience.

**Availability:** The source code for the latest version of easyClock is available at https://github.com/HungryFly/easyClock_for_CircadianRhythms. To download the installation package and access version updates, please refer to https://github.com/Dr-WUBINBIN/easyClock/releases.

## Introduction

The circadian clock governs many physiological processes and animal behaviors, ranging from gene expression to activity, sleep, and feeding^1^. Various tools have been used for acquiring behavioral and physiological data, such as the Activity Recording CAFE (ARC)^2^, *Drosophila* Activity Monitor (DAM)^3^, or multiomics^4^, which collectively generate substantial data. Numerous analysis methods can be employed to determine data rhythmicity, including JTK_CYCLE^5^, Cosinor analysis^6^, or RAIN^7^. However, most of these tools require some coding expertise^5–10^. Other graphical user interface (GUI) analyzers, such as ClockLab^11^ and ShinyR-DAM^12^, are either not freely accessible to users or attached to specific behaviors. Web-based analysis solutions reduce operational difficulties but pose the risk of inadequate maintenance. While recent circadian rhythm analyzers tend to simplify coding operations and optimize GUI design^13^, a more efficient and user-friendly tool would facilitate analysis of data collected from high-throughput experiments. To overcome operational barriers and enhance analysis efficiency, a GUI application named “easyClock” was developed to facilitate the local processing of multiple time series sets. The free-to-use easyClock application offers an intuitive interface and efficient, comprehensive mathematical methods, thereby improving user experience in data analysis and presentation within the field of chronobiology.

## Results

### Data Input and Preview

The easyClock package applies user-centric design principles, offering a streamlined, one-step installation process. Its clean, intuitive interface is organized into modular panels that guide users sequentially through each stage, beginning with data input (Figure 1A). As a lightweight tool, easyClock focuses on downstream analysis and visualization after basic pre-processing, rather than incorporating parsing pipelines for diverse data acquisition systems, ensuring adaptability across various experimental setups. Users can load up to three CSV files simultaneously, each with user-defined time intervals (Figures 1B & 2). This flexibility is especially valuable for comparing different phenotypes from the same groups, even when time intervals differ.

**Figure 1:**
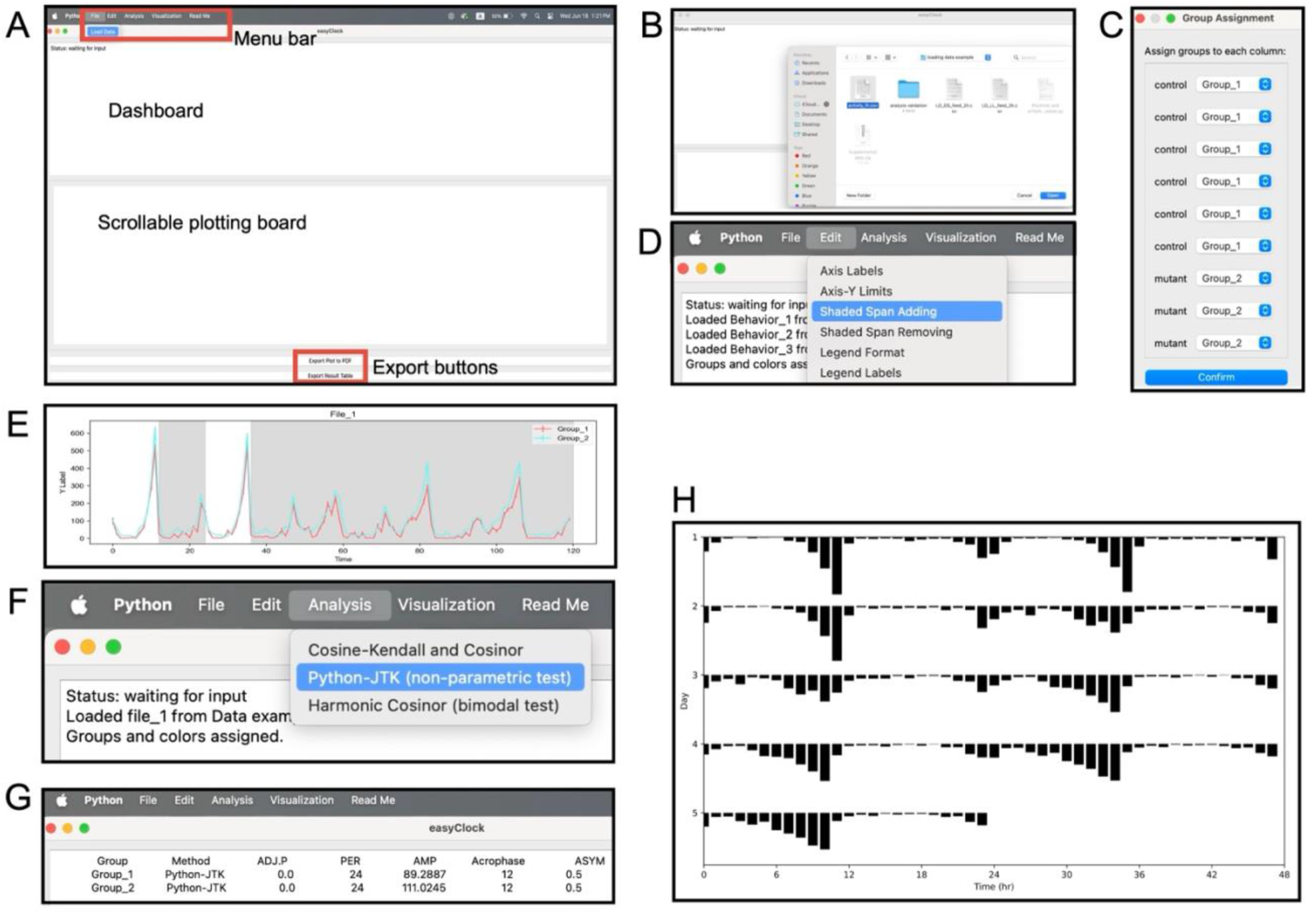
Interface of the easyClock application. (A) The easyClock interface is comprised of four distinct components: a menu bar, a dashboard, a scrollable plotting board, and export buttons. The menu bar, situated at the top of the interface, comprises four menus designed to facilitate file input (A, B, and C), plot editing (D and E), data analysis (F and G), and visualization (H) functionalities. Below the menu bar, the dashboard presents a comprehensive display of all execution information (F) and analysis outcomes (G). The scrollable plotting board serves as a visual representation of the data (A and E), offering a preview of all datasets and enabling real-time refreshes through edit capabilities. At the bottom, two buttons (A) are positioned to facilitate the export of plots and analysis results displayed on the boards. The analysis methods typically report four key parameters (G), including adjusted *p*-values (ADJ.P), period (PER), amplitude (AMP), and acrophase (peak time).

**Figure 2:**
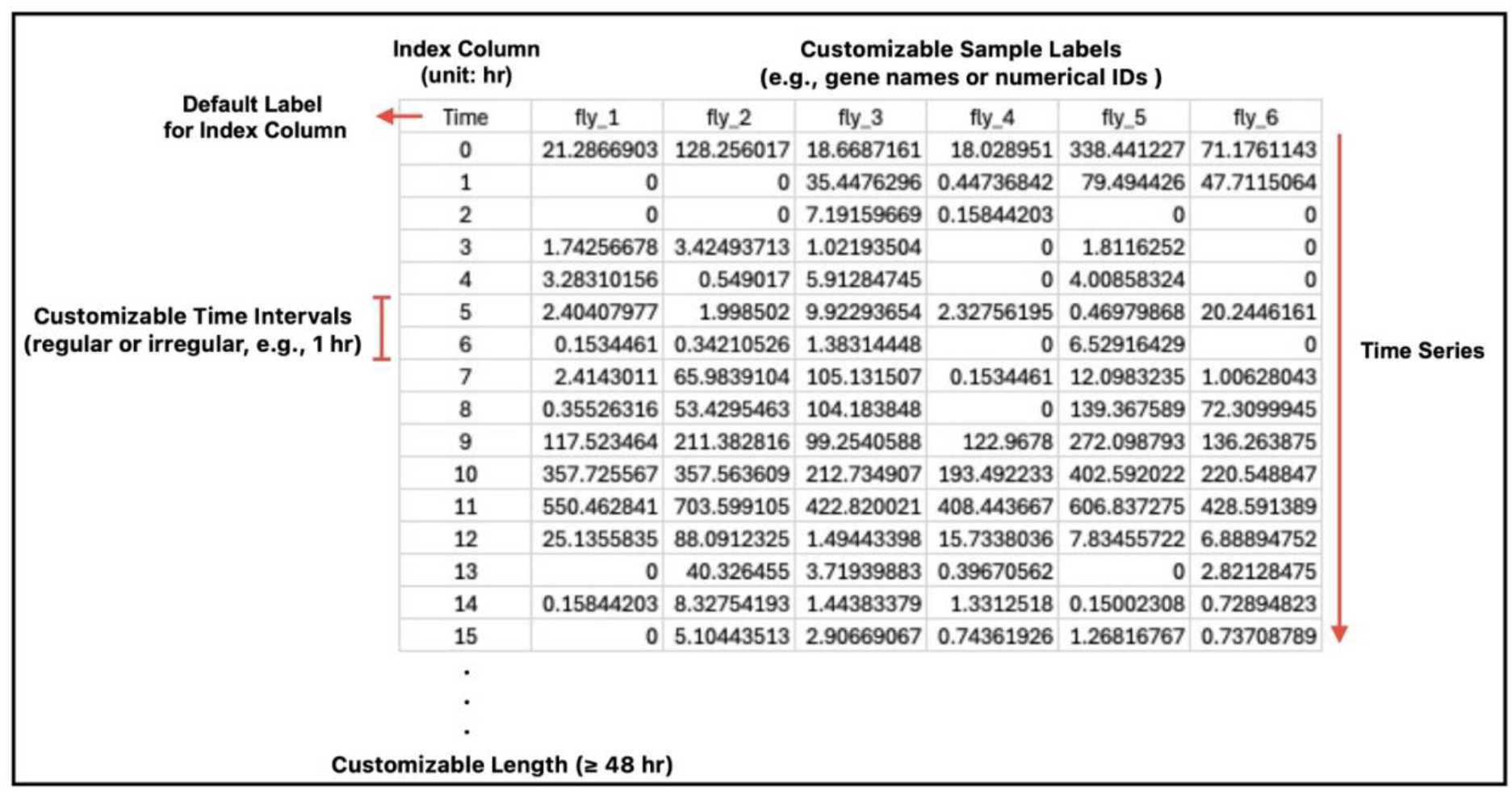
Example of input data format. The easyClock application allows users to load (or cancel) up to three CSV files, each occupying a separate slot. Each file must include an index column with a default label (“Time”), sample labels, and time-series data. The application supports both regular and irregular time intervals. Sample labels should be placed in the first row, starting from the second column, and can be customized as needed. Upon loading, group assignments are automatically generated from file #1 by grouping samples with identical labels; these groupings can be manually adjusted. When analyzing multiple files, groups in subsequent files are assigned automatically based on label matches with file #1. For robust and accurate analyses, at least 48 hours of data is recommended, although there is no minimum requirement for data resolution (i.e., the number of data points per sample within a given period).

Upon data loading, samples in file #1 that share the same label are automatically grouped. An initial dialog box allows users to customize these groups and assign colors (Figure 1C). Manual group assignment is primarily intended for single-file analyses, as subsequent files are grouped automatically based on label matches to file #1. The “Edit” menu (Figure 1D) provides a personalized preview of all datasets as conventional time-series plots (Figure 1E). A notable feature in this menu is the shaded span setting, which allows users to quickly add background colors to any data range to distinguish light/dark periods or other experimental conditions (Figures 1D-E). Data previews can be exported as publication-quality figures or used as references for subsequent analysis.

### Circadian Rhythm Analysis

Following data input and preview, users can select a file for analysis (Figure 1F & G). All time series within the selected file are analyzed, with users prompted to specify the period of interest; a minimum of 48 hours is recommended. The current version of easyClock includes four analytical methods: Cosinor, Cosine-Kendall, Python-JTK, and Harmonic-Cosinor. Both the Cosinor and Cosine-Kendall are designed for analyzing cosine-like rhythmic data.

In easyClock, the Cosinor method applies ordinary least squares regression to fit a linearized cosine model to the data^6^: *y* = *M* + β_1_cos(*wx*) + β_2_sin(*wx*) + ε. Rhythmicity is assessed using an *F*-test for the null hypothesis: β_1_ = β_2_ = 0. To control the false discovery rate across multiple comparisons, *p*-values are adjusted using the Benjamini-Hochberg procedure. The Cosine-Kendall method extends the Cosinor approach by ranking the data to reduce the influence of outliers. It measures concordance between the ranked input and each ranked cosine reference using Kendall’s tau^14^. The lowest *p*-value from the best-fitting model is selected, and Bonferroni correction is applied for multiple testing.

The easyClock package also features a non-parametric test, Python-JTK, which is a rewritten and modified implementation of the original JTK_CYCLE algorithm. Originally developed for R, JTK_CYCLE is widely used for detecting circadian rhythms^5^. It operates by performing rank-based comparisons between the input data and a library of discrete, triangle-shaped templates that may be either symmetric or asymmetric. In easyClock, Python-JTK employs Kendall’s tau to measure the similarity between the ranked input data and each template, replacing the Jonckheere–Terpstra (JT) test used in the original algorithm. A key advantage of Python-JTK in easyClock is its ability to automatically accommodate both regular and irregular time intervals, making it suitable for a wide range of experimental designs. To improve computational efficiency, easyClock allows users to customize parameters such as expected period, acrophase, and asymmetry, thereby reducing the number of template comparisons required and accelerating the analysis.

*Drosophila* behaviors, such as sleep and locomotor activity, frequently display bimodal patterns under 12-/12-hr light/dark cycles^15^. Traditional unimodal analysis methods are inadequate for capturing these complex rhythms. To address this, easyClock incorporates Harmonic-Cosinor analysis (also known as multiple-component cosinor^16^), which enables the fitting of bimodal waveforms by combining multiple sine and cosine components within each cycle (two components in the current version). The concordance between input data and each composite waveform reference is assessed using Kendall’s tau, chosen for its robustness, as demonstrated in the Cosinor-Kendall and Python-JTK methods.

Circadian rhythm data often display considerable variation in both oscillation amplitude and waveform shape. To evaluate the effectiveness of the four analysis methods in handling diverse data types, we simulated a series of arrhythmic and rhythmic datasets using Python (Sub-Table 1 & Supplemental Data). Among 50 groups of pseudo-randomly generated time series, Cosine-Kendall, Python-JTK, and Harmonic-Cosinor each achieved accuracy rates exceeding 98%, whereas the Cosinor method demonstrated comparatively lower accuracy in avoiding false positive detections of rhythmicity (Sub-Table 2). These results suggest that the use of Kendall’s tau effectively reduces the false positive rate.

**Sub-Table 1:**
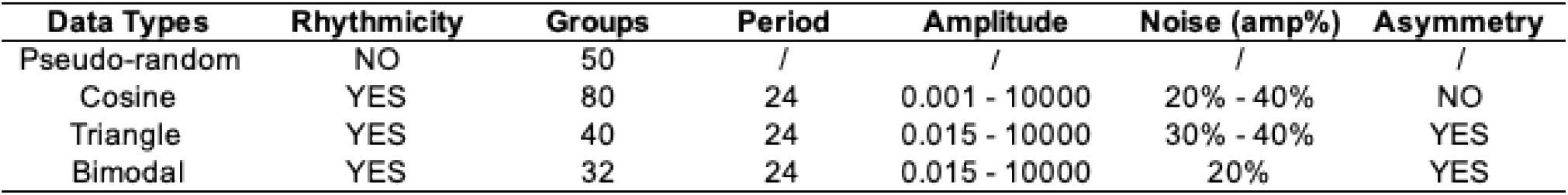
Data simulation.

**Sub-Table 2:** Data detection.

| Methods | Pseudo-random | Cosine | Triangle | Bimodal |
| --- | --- | --- | --- | --- |
| Cosinor | 46/50 | 80/80 | 40/40 | 32/32 |
| Cosinor-Kendall | 50/50 | 80/80 | High FNR | High FNR |
| Python-JTK | 49/50 | 80/80 | 40/40 | 32/32 |
| Harmonic-Cosinor | 49/50 | 80/80 | High FNR | 32/32 |

To further assess method performance, we simulated various types of circadian rhythm data—including single-component cosine, asymmetric triangle waves, and bimodal patterns (Sub-Table 1 & Supplemental Data). All three unimodal methods excelled at detecting cosine waveforms, even under conditions of low amplitude and high noise (Sub-Table 2 & 3). For asymmetric triangle waves, Python-JTK outperformed the other methods, especially with highly asymmetric signals (Sub-Table 2 & 3). In the case of bimodal data, Harmonic-Cosinor provided superior accuracy in period detection compared to unimodal approaches (Sub-Table 3).

**Sub-Table 3:** Period deviation and method recommendation.

| Methods | FPR | Cosine | Triangle | Bimodal |
| --- | --- | --- | --- | --- |
| Cosinor | 4/50 | 0 - 1.5 ☆ | 0 - 4.0 | 0 - 3.5 |
| Cosinor-Kendall | 0/50 | 0 - 2.0 ☆☆ | / | / |
| Python-JTK | 1/50 | 0 - 1.0 ☆☆☆ | 0 - 2.0 ☆☆☆ | 0 - 2.0 ☆☆ |
| Harmonic-Cosinor | 1/50 | 0 - 3.5 | / | 0 - 0.5 ☆☆☆ |

### Visualization

Actograms are widely used in circadian rhythm research for visualizing rhythmic patterns and phase shifts in time-series data. In addition to data preview, easyClock enables automatic generation of actograms (Figure 1H). Users can quickly export double-plotted actograms for any selected group with a few intuitive clicks, without the need to adjust parameters.

Furthermore, easyClock provides visualization of the fitting models following analysis. For each dataset, it displays the best fit of the selected fitting model on the original or group-averaged time series, allowing users to visually assess the quality of rhythmicity detection. Depending on the selected method, users can visualize the fitting results of Cosinor analysis (Figure 3A), Cosine-Kendall (Figure 3B), Python-JTK (Figure 3C), and Harmonic-Cosinor (Figure 3D).

**Figure 3:**
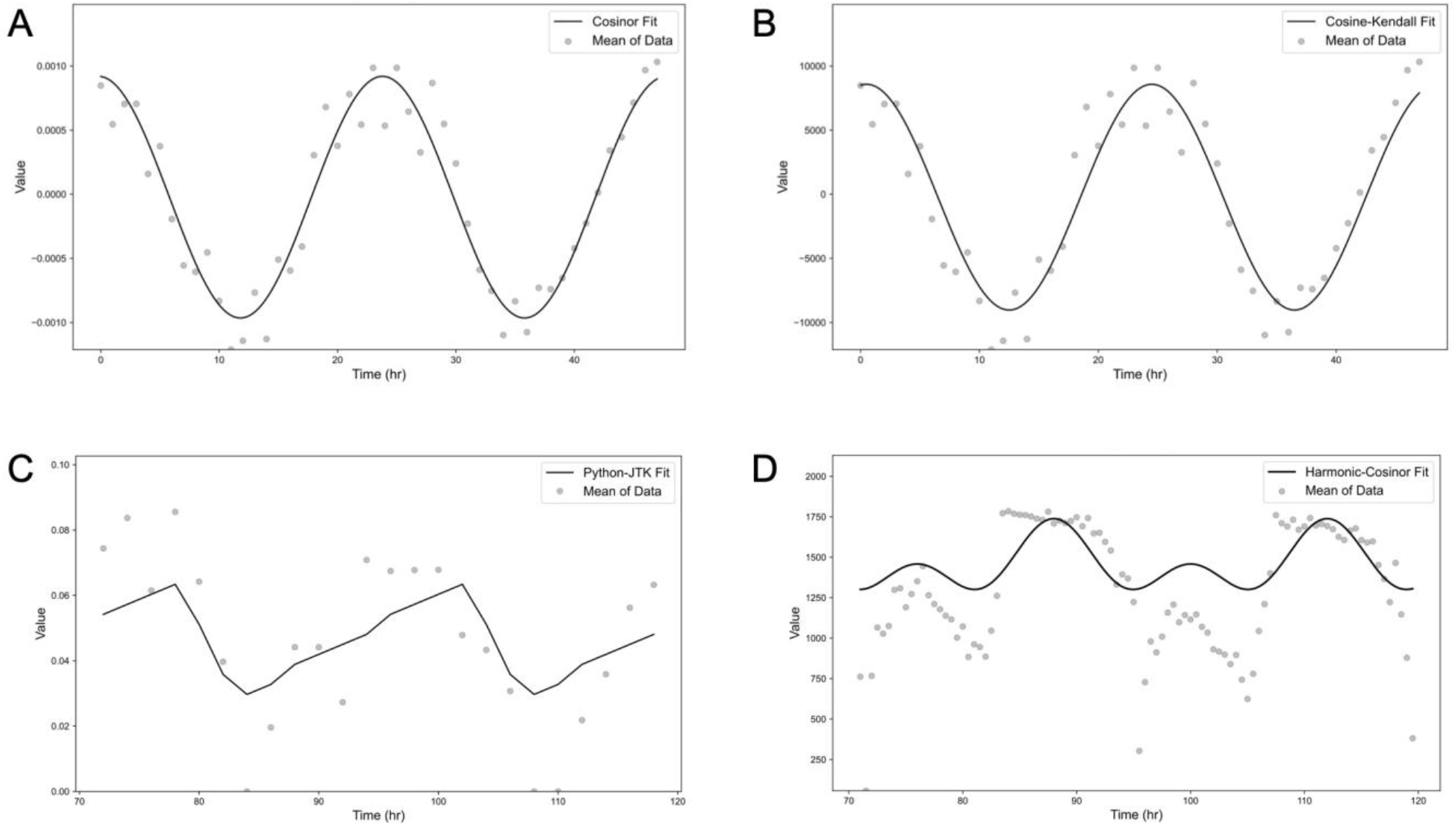
Visualization of model fittings across diverse waveforms. easyClock applies appropriate analysis methods to different data waveforms, enabling visualization of the fitting models alongside the individual or group-averaged time series. Representative examples of model fits include: (A) Cosinor, (B) Cosinor-Kendall, (C) Python-JTK, and (D) Harmonic-Cosinor.

## Conclusion

Over the past several decades, various analytical and statistical methods have been developed and adopted in chronobiology, ranging from visual inspection of actograms to sophisticated mathematical statistics^17^. Among these, Analysis of Variance (ANOVA) has served as a fundamental tool for detecting the oscillatory characteristics of circadian data. However, ANOVA alone cannot directly demonstrate the presence of rhythmicity. To overcome this limitation, model-fitting techniques have been introduced. In response to the need for a versatile, accessible, and user-friendly circadian analysis toolkit, we developed easyClock—a GUI application that integrates comprehensive fitting models for circadian rhythm detection.

It is important to note that the current analytical methods implemented in easyClock operate on averaged time-series data, either across groups or for individual time series. While this approach enables rapid identification of rhythmic patterns, it does not fully capture within-group variability. Future iterations of easyClock may incorporate statistical models that account for inter-individual variance, thereby providing a more robust analysis of circadian rhythms.

In easyClock, amplitude serves as a primary measure of rhythmic strength—a key concept in chronobiology that distinguishes robust circadian patterns from weak or noisy signals. Weak rhythms are typically characterized by irregular peaks, inconsistent phases, or diminished amplitudes. In addition to amplitude-based assessments, the chi-square periodogram (CSP) is commonly employed to quantify rhythmic strength by evaluating deviations from the global mean at each circadian time point^18^. Although high CSP values generally indicate strong rhythmicity, this method can be susceptible to false positives in the presence of substantial fluctuations within the time series^19^. As an extensible platform, future updates of easyClock intend to integrate more comprehensive and reliable methods for assessing rhythmic strength.

## Acknowledgments

We thank Dr. Yutong Xiao (Max Planck Florida Institute for Neuroscience), Alayna Garland (Kenan Fellow), and Dr. Qiankun He (Zhengzhou University) for participating in the beta-testing of this desktop application and providing valuable feedback. This work was funded by the NIH (R01DC020031, W.W.J.).

## Author Contributions

Conceptualization, algorithm development, and software implementation: B.W.; Supervision: W.W.J.; Writing—original draft: B.W.; Writing—review and editing: B.W. and W.W.J.

## Table: Comparison of rhythmicity detection methods in easyClock

To evaluate the performance of the analysis methods in easyClock, various types of 48-hr time series were simulated using Python (Sub-Table 1). Arrhythmic datasets (50 groups) were generated pseudo-randomly, while rhythmic datasets featured cosinor, triangular, or bimodal waveforms with varying amplitudes (amp) and asymmetries. Additional pseudo-random noise was added to the data as a percentage of amplitude. Four analysis methods were applied to detect rhythmicity in these datasets (Sub-Table 2). Each fraction represents the proportion of groups correctly identified as arrhythmic (in pseudo-random data) or rhythmic (in circadian datasets) by each method. “High FNR” indicates a high false negative rate (>35%) for a given method when analyzing incompatible data types. The methods displayed differing levels of effectiveness depending on the waveform type (Sub-Table 3). The false positive rate (FPR) was calculated based on results from Sub-Table 2. Period deviation is reported as a range (e.g., 0-1.5), reflecting the extent to which the estimated period differed from the expected value across tested groups. Recommendation symbols (☆ to ☆☆☆) indicate increasing levels of suitability based on each method’s ability to detect periods accurately and minimize FPR.

## References

1. Panda, S., Hogenesch, J. B. & Kay, S. A. Circadian rhythms from flies to human. Nature 417, 329–335 (2002).

2. Murphy, K. R., Park, J. H., Huber, R. & Ja, W. W. Simultaneous measurement of sleep and feeding in individual Drosophila. Nat Protoc 12, 2355–2359 (2017).

3. Pfeiffenberger, C., Lear, B. C., Keegan, K. P. & Allada, R. Locomotor Activity Level Monitoring Using the Drosophila Activity Monitoring (DAM) System: Figure 1. Cold Spring Harb Protoc 2010, pdb.prot5518 (2010).

4. Huang, R. et al. Multi-omics profiling reveals rhythmic liver function shaped by meal timing. Nat Commun 14, (2023).

5. Hughes, M. E., Hogenesch, J. B. & Kornacker, K. JTK_CYCLE: An Efficient Nonparametric Algorithm for Detecting Rhythmic Components in Genome-Scale Data Sets. J Biol Rhythms 25, 372–380 (2010).

6. Moškon, M. CosinorPy: a python package for cosinor-based rhythmometry. BMC Bioinformatics 21, 485 (2020).

7. Thaben, P. F. & Westermark, P. O. Detecting Rhythms in Time Series with RAIN. J Biol Rhythms 29, 391–400 (2014).

8. Geissmann, Q., Garcia Rodriguez, L., Beckwith, E. J. & Gilestro, G. F. Rethomics: An R framework to analyse high-throughput behavioural data. PLoS ONE 14, e0209331 (2019).

9. Singer, J. M. & Hughey, J. J. LimoRhyde: A Flexible Approach for Differential Analysis of Rhythmic Transcriptome Data. J Biol Rhythms 34, 5–18 (2019).

10. Wu, G., Anafi, R. C., Hughes, M. E., Kornacker, K. & Hogenesch, J. B. MetaCycle: an integrated R package to evaluate periodicity in large scale data. Bioinformatics 32, 3351–3353 (2016).

11. Brenna, A., Ripperger, J. & Albrecht, U. Locomotor Activity Monitoring in Mice to Study the Phase Shift of Circadian Rhythms Using ClockLab (Actimetrics). BIO-PROTOCOL 15, (2025).

12. Cichewicz, K. & Hirsh, J. ShinyR-DAM: a program analyzing Drosophila activity, sleep and circadian rhythms. Commun Biol 1, 25 (2018).

13. Abhilash, L. & Sheeba, V. RhythmicAlly: Your R and Shiny–Based Open-Source Ally for the Analysis of Biological Rhythms. J Biol Rhythms 34, 551–561 (2019).

14. Hamed, K. H. The distribution of Kendall’s tau for testing the significance of cross-correlation in persistent data. Hydrological Sciences Journal 56, 841–853 (2011).

15. Xia, X. et al. Regulation of circadian rhythm and sleep by miR-375-timeless interaction in Drosophila. FASEB j. 34, 16536–16551 (2020).

16. Cornelissen, G. Cosinor-based rhythmometry. Theor Biol Med Model 11, 16 (2014).

17. Refinetti, R., Cornélissen, G. & Halberg, F. Procedures for numerical analysis of circadian rhythms. Biological Rhythm Research 38, 275–325 (2007).

18. Sokolove, P. G. & Bushell, W. N. The chi square periodogram: Its utility for analysis of circadian rhythms. Journal of Theoretical Biology 72, 131–160 (1978).

19. Tackenberg, M. C. & Hughey, J. J. The risks of using the chi-square periodogram to estimate the period of biological rhythms. PLoS Comput Biol 17, e1008567 (2021).

